# A novel algorithm to accurately classify metagenomic sequences

**DOI:** 10.1101/2020.10.01.321067

**Authors:** Subrata Saha, Zigeng Wang, Sanguthevar Rajasekaran

## Abstract

Widespread availability of next-generation sequencing (NGS) technologies has prompted a recent surge in interest in the microbiome. As a consequence, metagenomics is a fast growing field in bioinformatics and computational biology. An important problem in analyzing metagenomic sequenced data is to identify the microbes present in the sample and figure out their relative abundances. In this article we propose a highly efficient algorithm dubbed as “Hybrid Metagenomic Sequence Classifier” (HMSC) to accurately detect microbes and their relative abundances in a metagenomic sample. The algorithmic approach is fundamentally different from other state-of-the-art algorithms currently existing in this domain. HMSC judiciously exploits both alignment-free and alignment-based approaches to accurately characterize metagenomic sequenced data. To demonstrate the effectiveness of HMSC we used 8 metagenomic sequencing datasets (2 *mock* and 6 *in silico* bacterial communities) produced by 3 different sequencing technologies (e.g., HiSeq, MiSeq, and NovaSeq) with realistic error models and abundance distribution. Rigorous experimental evaluations show that HMSC is indeed an effective, scalable, and efficient algorithm compared to the other state-of-the-art methods in terms of accuracy, memory, and runtime.

**Availability of data and materials:** The implementations and the datasets we used are freely available for non-commercial purposes. They can be downloaded from: https://drive.google.com/drive/folders/132k5E5xqpkw7olFjzYwjWNjyHFrqJITe?usp=sharing

## 1 Introduction

Although we are normally unable to see microbes, they run the world. Microbes are indispensable for every part of our human life - truly speaking - all life on Earth! They influence our daily life in a myriad ways. For an example, the microbes living in the human gut and mouth enable us to extract energy from food. We could not be able to digest food without them. Microbes also insulate us against disease-causing agents. Metagenomics is a strong tool that can be used to decipher microbial communities by directly sampling genetic materials from their natural habitats. It can be directly applicable a wide variety of domains to solve practical challenges such as biofuels, biotechnology, food safety, agriculture, medications, etc. Metagenomic sequencing shows promise as a sensitive and rapid method to diagnose infectious diseases by comparing genetic material found in a patient’s sample to a database of thousands of bacteria, viruses, and other pathogens. There is also strong evidence that microbes may contribute to many non–infectious chronic diseases such as some forms of cancer, coronary heart disease, and diabetes. It offers to decipher the role of the human microbiome (the collective genome of our symbionts) in health and disease in individuals and populations, and the development of novel diagnostic and treatment strategies based on this knowledge. In short, it has the capability to impact the humanity indeed!

Classifying metagenomic sequences in the metagenomic sample is a very challenging task due to these facts: (1) researchers have sequenced complete genomes of thousands of microbes. The total size of the genomes of such microbes is hundreds of gigabytes. Again, metagenomic sample can contain millions to billions of biological sequencing reads. The total size of these reads can range from gigabytes to terabytes. To detect the microbes in the sample, we must align the metagenomic reads onto the known reference genomes of microbes. Processing such a huge data is very difficult to accomplish as we have time and memory bound; and (2) different microbes can contain similar genomic regions in their genomes. Reads in the given metagenomic sample may refer to different microbes that are not present in the sample. Consequently, there might be a lot of false identification if we just align the reads onto the references.

Several subsequence-based approaches exist in this domain to determine the identity and relative abundance of microbes in a heterogenous microbial sample. These methods suffer from low classification accuracy, high execution time, and high memory usage. The users also need to post-process the output. Thus, we need efficient and effective computational technique to quickly and accurately identify microbes present in metagenomic sample. To address the issues stated above we developed a highly efficient method to correctly identify and estimate the abundance of microbes in the metagenomic sample. To process the huge amount of data and correctly identify microbes we have come up with a very novel computational method. At first, we reduce the size of a genome sequence of a known microbe by extracting only the unique regions of that genome. These unique regions are not present in any other microbes. To do this efficiently we collect a set of subsequences from each of the reference genomes. These subsequences are unique among all the reference genomes. The number of such subsequences could be hundreds of millions. To process such number of subsequences efficiently in limited resources, we offer a novel out-of-core algorithm. These unique subsequences are used to extract discriminating regions across a genome. Those regions are then used to build a model sequence of a genome. Instead of using the full genome sequences, these pre-built model sequences are then employed to accurately profile all taxonomic levels and relative abundances of the microbes present in a metagenomic sample. Rigorous experimental evaluations show that our algorithm HMSC is highly efficient in terms of accuracy, execution time, and memory footprint.

## 2 Methods

### 2.1 Approach

The traditional approach to solve the metagenomic sequence classification is to align each input read to a large collection of reference genomes using alignment software such as BLAST [3] or MegaBlast [17]. However, aligning the reads to the reference sequences requires a huge computing time. One way of salvaging execution time could be to align the reads to a select marker genes in the reference genome instead of the whole genome. This approach is followed in Metaphan [25], Metaphyler [13], Motu [23] and Megan [9]. However, this approach also becomes infeasible when there are more and more of reference genomes and the total number of reads grows more and more.

Researchers have tried several alignment free methods. The most popular among such methods is based on *k*-mer spectrum analysis which uses a database of distinct subsequences of length *k*, or in short *k*-mers, from the reference sequence to classify the reads. If a read has distinct *k*-mers from multiple reference genomes then it is assumed to be from their lowest common taxonomic ancestor (see e.g., [26]). The algorithms using this broad approach differ in the way the database is built and queried. LMAT [1] is one of the first such algorithms. Subsequent improvements are: Kraken [26] which improves the speed and memory usage by employing a classification tree, Clark [20] improves memory usage further by storing only a reduced set of target specific *k*-mers. Clark-S [19] improves the specificity of CLARK by sacrificing a little speed and memory. Metacache [18] improves the memory usage further by a novel application of minhashing technique on a subset of the context aware *k*-mers to reduce memory usage. MetaOthello [14] uses a probabilistic hashing classifier and improves the memory usage and k-mer query efficiency with a novel *I* -Othello data structure. LiveKraken [24] classifies reads in realtime from raw data streams from Illumina sequencers. KrakenUniq [2] efficiently assesses the coverage of unique k-mers in each species and gives better recall and precision.

Kaiju [16] uses the same *k*-mer based approach, however, exploits the fact that microbial and viral genomes are typically densely packed with protein-coding genes which are more conserved and more tolerant to sequencing errors because of the degeneracy of the genetic code. Some applications require an estimate of the relative abundance of constituent species. Using a probabilistic method based on Byesian likelihood, Braken [15] augments the output of Kraken with estimated abundance. There have been few alignment free methods other than the *k*-mer spectrum analysis as well. MetaKallisto [22] uses pseudo alignments. WGSQuikr [12] and Metapallette [11] use a compressed sensing approach where the abundance is estimated by solving a linear system of equations on the *k*-mer spectrum profile of input reads and that of the reference sequences. TaxMap [5] uses a novel database compression algorithm to store the LCA information and achieves better precision and sensitivity.

Recently, deep learning models [7], such as convolutional neural network (CNN) and deep belief network, have been tested on a small subset of bacteria taxonomic classification, but it is still challenging for the classification for the entire bacteria domain.

### 2.2 Our algorithms

There are 3 major steps in our proposed algorithm HMSC. The first step involves collecting a set of unique *k*-mers from each of the genomic sequences of interest. This step is different from the *k*-mer counting problem. In *k*-mer counting we compile all the distinct *k*-mers present in the input sequences together with the frequency of each *k*-mer. A *k*-mer can be found in more than one genomic sequence. Conversely, each unique *k*-mer can be found in one and only one genome sequence. The second step involves finding a set of *discriminating regions* from each genome. We then build a model sequence for each genome by adopting the discriminating regions. Instead of using the full genome sequences, these pre-built model sequences are then employed to accurately profile all the 8 taxonomic ranks. We also estimate approximate abundance of all the 8 taxonomic ranks residing in a given metagenomic sample. Experimental evaluations show that HMSC is highly efficient in terms of accuracy, execution time and memory footprint. Next we describe the algorithmic steps of HMSC in detail.

#### 2.2.1 Unique k-mer mining

At the beginning we identify a set of unique *k*-mers from the given set of target genomes *G* = {*g*_1_, *g*_2_, *g*_3_, *…, g*_*n*_}. A *k*-mer is said to be unique if it (and its reverse complement) occurs in only one of the genomes *g*_*i*_ ∈ *G* where 1 ≤ *i* ≤ *n*. However, if we want to search for unique *k*-mers from the set of all such *k*-mers in one single pass, it will be a very memory intensive and time consuming procedure. In our proposed algorithm we employ around 6*k* bacterial genomes and each genome contains nearly 4.5*M* nucleotides on an average! To reduce the memory footprint we partition *G* into *P* non-overlapping parts. In each such part *p*_*j*_ (where 1 ≤ *j* ≤ *P*) we search for *k*-mers that are unique across all the genomes. To reduce the search space we randomly select a subset of all the *k*-mers present in the genomic sequences.

For each partition *p*_*j*_ ∈ *P* we maintain two data structures to find the unique *k*-mers in each genome. Hash table *H* is used to record unique *k*-mers. Each key of the hash table *H* represents a *k*-mer and its corresponding value refers to the associated genome id and start index. Hash set *S* contains the non-unique *k*-mers found in more than one genome. In both data structures the expected time complexity to search, update, or delete operations is *O*(1). After collecting locally unique *k*-mers by utilizing *H* and *S* for each genome in one part, we collect the unique set of *k*-mers from the reduced set of locally unique *k*-mers found in the previous step by checking if it occurs in the other genomes. We save the unique *k*-mers picked from each part with the associated genome ids and start indices in the disk. Next we describe the proposed method in detail.

For each genome *g* in a part *p*_*j*_, 1 ≤ *j* ≤ *P*, we process it as follows. We pick a random subset of *k*-mers in *g*. Let *x′* and *x″* be any such *k*-mer and its reverse complement, respectively. Suppose *x* is the lexicographically smallest of *x′* and *x″*. We first check if *x* is already declared as non-unique by searching for it in the hash set *S*. If *S* already contains *x*, it is non-unique. Otherwise we search for *x* in the hash table *H*. There are two possibilities: there is an entry for *x* in *H* or there is no entry for *x* in *H*. If there is an entry for *x* in *H*, there are two possibilities: (1) The *k*-mer in the entry corresponds to another genome or (2) It corresponds to the same genome (as that of *x*). In the former case we declare *x* as non-unique by recording *x* in *S* and delete the entry from *H*. If the later is the case, we do not do anything. If we do not find any entry for *x* in *H*, we create an entry for *x* in *H* associated with its genome id and position. At the end of processing all the randomly picked *k*-mers in all the genomes in *p*_*j*_, in the above manner, *H* has all the locally unique *k*-mers in *p*_*j*_. Please, note that these locally unique *k*-mers are only unique with respect to the randomly picked *k*-mers from the genomes in *p*_*j*_. From out of these locally unique *k*-mers, we identify globally unique *k*-mers. To find the globally unique *k*-mers, we iteratively retrieve each genome *g′* ∈ *G* from the disk. To check the duplicity efficiently we build a hash set *S′* containing the lexicographically smallest *k*-mers of *g′*. I.e., for each *k*-mer in the retrieved genome *g′*, we insert into *S′* the lexicographically smallest one between it and its reverse complement. Now the locally unique set of *k*-mers are checked for duplicity against *S′*. Please, note that the genomes are retrieved from the disk into the main memory one at a time. The locally unique *k*-mers will also be checked for duplicity with respect to the genomes in *p*_*j*_. Once we identify the globally unique *k*-mers in *p*_*j*_, we save them together with their positions and genome ids in the disk.

As noted earlier, we partition *G* into *P* parts (for some suitable value of *P*) to reduce the memory footprint. For each of the *P* parts we follow the same procedure as described above. We save the globally unique *k*-mers along with their genome ids and positions in the disk for each part. In the experimentation we set *k* = 31. Details of the algorithm can be found in Algorithm 1.

#### 2.2.2 Model sequence formation

In this step we build a model sequence for each genome *g* ∈ *G*. Let the set of unique *k*-mers in *g* (found in the previous step) be *u*. At first we sort *u* in increasing order with respect to the starting coordinates of the *k*-mers. Next we cluster the sorted *k*-mers in such a way that in any cluster the distance between any pair of consecutive *k*-mers is ≤ a threshold *t*_1_. Let a pair of consecutive *k*-mers be *k′* and *k″*. Then the distance between the end position of *k′* and the start position of *k″* will be no more than *t*_1_. We linearly search through the sorted list of *k*-mers and build a new cluster when the distance between any pair of consecutive *k*-mers is *> t*_1_. Let the set of clusters be *C*. Clearly, each cluster *c* ∈ *C* contains a set of consecutive *k*-mers where the distance between any two consecutive *k*-mers is ≤ *t*_1_. Consequently, each cluster *c* ∈ *C* contains a discriminating region of the genome *g*. We extract a region from *g* by using the start index of the first *k*-mer and the end index of the last *k*-mer in *c*. We call such a region *discriminating* since if any read *r* having length ≥ *t*_1_ + 2*k* is aligned onto such a region, then *r* will fully contain at least 1 unique *k*-mer. In our experiment we set *t*_1_ = 40.

We sort all such regions from all the clusters *c* based on decreasing order of their lengths. We discard some regions having length ≤ *t*_2_, a user defined threshold. We append a special string of length 4 containing a special character “#” at the end of each region. We concatenate all such regions of a genome *g* to build a model sequence. Because of this special string no aligned read will contain the junction of any 2 regions given that the mismatch threshold *d <* 4. Let these model sequences are *m*_1_, *m*_2_, *…, m*_*n*_ where *m*_*i*_ is the model sequence of *g*_*i*_, 1 ≤ *i* ≤ *n*. In our experiment we set *t*_2_ = 300. Details of the algorithm can be found in Algorithm 2.

#### 2.2.3 Taxonomic ranks identification

In this step we are inferring all the 8 taxonomic ranks (e.g., subspecies, species, genus, family, order, class, phylum, and kingdom) and their corresponding relative abundances from a given metagenomic sample *V*. Metagenomic sequencing reads contained in *V* are aligned onto each of the model sequences *m*_*i*_ built in the previous step. Suppose a read *r* ∈ *V* is aligned onto a model sequence *m*_*i*_ within a certain mismatch threshold *d* (we set *d* = 0 in our experiment). We can say that the read *r* belongs to the genome *g*_*i*_ with a high level of confidence. This is referred to as a *hit*.

If a read *r* is aligned onto multiple model sequences *m*_*i*_ from multiple genomes *g*_*i*_ ∈ *G*, then the read *r* is assigned to all of those genomes *g*_*i*_. Since the taxonomic profiling of HMSC is based on the model sequences of the genomes, not all the reads *r* from the metagenomic sample *V* will be classified. This is due to the fact that the model sequences may not contain all the stretches of the original genomes. We estimate the abundance of a specific taxonomic rank using the number of hits. Consider a specific genome *g*_*i*_. Let the taxonomic rank of *g*_*i*_ be *t*_*i*_. We estimate *t*_*i*_ by taking the ratio of the hits in *m*_*i*_ with respect to that taxonomic rank to the total number of hits (across all the model sequences).

The accuracy of our algorithm has been measured using the ground truth. For instance, if we employ synthetic data, then we will know what species are represented in the sample *V* and also the number of reads in *V* corresponding to each of the species.

### 2.3 Analyses

#### 2.3.1 Number of possible unique *k*-mers

In this section we compute the expected number of unique *k*-mers under a random model. This number will be essential to estimate the run time of our algorithm. Let a database consist of *n* genomes of length *m* each. Consider a random model where each character of each genome is uniformly randomly picked from the alphabet {*g, c, t, a*}. Let the genomes in the datatbase be *g*_1_, *g*_2_, *…, g*_*n*_. Let *x* be any *k*-mer of *g*_*i*_ (for any *i*). If *y* is any *k*-mer of *g*_*j*_ (with *i* = *j*), the probability that *x* = *y* is 4^*−k*^. This in turn means that the probability that *x* matches any *k*-mer in any other genome is ≤ (*n* − 1)(*m* − *k* + 1)4^*−k*^. As a result, the probability that *x* is unique, i.e., it does not occur in any other genome is ≥ 1 − (*n* − 1)(*m* − *k* + 1)4^*−k*^. This probability could be very high. For example, if *k* = 40, *m* = 100, 000, and *n* = 200, the above probability is around 1 − 1.645 × 10^*−*17^. This implies that when *k* is large, each of the *k*-mers is likely to be unique.

We can compute the expected number of matches as follows. As stated before, the probability that *x* = *y* is 4^*−k*^. Thus the expected number of *k*-mers of *g*_*j*_ that equal *x* is ≤ 4^*−k*^ (*m* − *k* + 1). Also, the expected number of *k*-mers from the genomes other than *g*_*i*_ that equal *x* is ≤ 4^*−k*^ (*n* − 1)(*m* − *k* + 1). When *k* = 40, *m* = 100, 000, and *n* = 200, this number is ≤ 1.645 × 10^*−*17^! We also note that the expected number of matching *k*-mer pairs is no more than 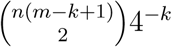. For the above example, this number is no more than 1.7 × 10^*−*10^.

From out of all the *n* genomic sequences, there are a total of *n*(*m* − *k* + 1) *k*-mers. It follows that the expected number of unique *k*-mers is ≥ *n*(*m* − *k* + 1) [1 − (*n* − 1)(*m* − *k* + 1)4^*−k*^] ≈ *n*(*m* − *k* + 1). This could indeed be very large. For this reason, we only employ a subset of these. We first find locally unique *k*-mers and from these we identify globally unique *k*-mers. To find locally unique *k*-mers, we only use randomly picked *k*-mers. Locally unique *k*-mers can also be found as follows. We use all the *k*-mers from all the genomes in one part to find unique *k*-mers. From these we find globally unique *k*-mers and from the globally unique *k*-mers, we pick a random subset.

Both the algorithms are equivalent (with different sampling rates). As a result, consider the second algorithm. If *s* is the sampling rate, then the expected number of unique *k*-mers picked will be *s n*(*m* − *k* + 1). Let *R* be any read in the metagenomic sample whose length is *r*. This read has (*r* − *k* + 1) *k*-mers. An important question is how many of the unique *k*-mers that we pick can be found in *R* by random chance. Let *x* be any unique *k*-mer that we pick and let *y* be any *k*-mer of *R*. Probability that *x* = *y* is 4^*−k*^. This means that the expected number of *k*-mers of *R* that will match *x* is ≤ 4^*−k*^ (*r* − *k* + 1). Also, the expected number of unique *k*-mers picked that will match any *k*-mer of *R* is ≤ *s n*(*m* − *k* + 1)4^*k*^ (*r* − *k* + 1). For example, if *k* = 40, *m* = 100, 000, *n* = 200, *r* = 100, and *s* = 0.1, then this number will be ≤ 10^*−*16^. This analysis ensures that the probability that a unique *k*-mer occurs in a read by random chance is very low.

#### 2.3.2 Time complexity - In-core model

In this section we estimate the run time of our algorithm in the case where all the input data can be stored in the main memory of the computer used. We also assume that we use the second algorithm for finding globally unique *k*-mers. Clearly, the run time of this algorithm will be an upper bound for the first algorithm’s run time. Let *n* and *m* denote the number of genomes and the length of each genome, respectively. Assume that *k* = *O*(1). The total number of *k*-mers in the database is *O*(*mn*).

##### Identification of unique k-mers

We spend an expected *O*(1) time for each operation on the hash tables *S* and *H*. For each *k*-mer *x* from each genome, in the worst case we perform a constant number of operations in *S* and *H*. As a result, the total expected time spent for each *k*-mer is *O*(1). Therefore, the expected time spent in identifying the unique *k*-mers is *O*(*mn*). Let *s* be the random sampling rate for picking the unique *k*-mers, i.e., we randomly pick a fraction *s* of all possible unique *k*-mers. Total expected time spent in this step is *O*(*mn*).

##### Model sequences formation

Consider any genome *g*_*i*_ (1 ≤ *i* ≤ *n*). Let *u*_*i*_ be the set of unique *k*-mers picked from *g*_*i*_. As shown in the above analysis, the expected number of *k*-mers in *u*_*i*_ is no more than *s n*_*i*_ where *n*_*i*_ is the length of *g*_*i*_. We sort these *k*-mers based on their starting indices. This can be done using the radix sort (or the bucket sort) algorithm in *O*(*s n*_*i*_) time. Followed by this sorting step, we can form the clusters by scanning through the sorted list once. This will take *n*_*i*_ time. From the clusters we obtain the discriminating regions. Subsequently, we sort the clusters based on their lengths. This will take *O*(|*C*|) time which is *O*(*s n*_*i*_). The total expected time spent thus far on *g*_*i*_ is *O*(*s n*_*i*_). Summing this over all the genomes, the total expected time is *O*(*s N*) where 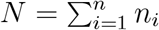.

The discriminating regions of *g*_*i*_ are then concatenated to get the model sequence *m*_*i*_ (for each *i*, 1 ≤ *i* ≤ *n*). Note that the length of the model sequence *g*_*i*_ can be as large as *n*_*i*_. In practice, it is much less than *n*_*i*_. The worst case time to form the model sequences is *O*(*mn*).

##### Taxonomic ranks computation

In this step we align all the reads in the sample *V* onto all the model sequences. Let *Q* be the number of reads and the length of each read be *r*. We use Bowtie to do the alignment. Bowtie will first construct a database using all the model sequences. The total length of all the model sequences is *O*(*mn*). Thus the time to construct the database is *O*(*mn*). Followed by this, each read will be aligned. The time spent for each read will be *O*(*r* + *h*) where *h* is the number of hits for the read. If we assume that *h* = *O*(1), then the total alignment time is *O*(*mn* + *Qr*).

In summary, the run time of the entire algorithm is *O*(*mn*+*Qr*) which is, clearly, asymptotically optimal.

#### 2.3.3 Time complexity - Out-of-core model

Now consider the out-of-core model. Assume that the core memory is not large enough to hold all the geomes. In fact this is the model that holds in practice and the one that we have implemented. Assume that the core memory can only hold 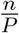 of the genomes. Since the time taken for the I/O operations is typically much more than the time spent on local computations, it is customary in the literature to report only the I/O complexity for out-of-core algorithms. We follow this custom for our MSC algorithm also.

##### Identification of unique k-mers

We bring 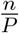 genomes at a time into the core memory and identify the unique *k*-mers in each part. Over all the parts, we do one pass through the entire genome dataset. Followed by this we bring all the unique *k*-mers from the disk into the core memory. Now we have to check which among these are indeed unique across all the genomes. This will take one more pass through the genome dataset. We assume that the randomly picked set of unique *k*-mers is small enough to fit in the core memory.

##### Building model sequences

Consider any genome *g*_*i*_ (1 ≤ *i* ≤ *n*). Let *u*_*i*_ be the set of unique *k*-mers picked from *g*_*i*_. As shown in the above analysis, the expected number of *k*-mers in *u*_*i*_ is no more than *s n*_*i*_ where *n*_*i*_ is the length of *g*_*i*_. We have to sort these *k*-mers based on their starting indices. Followed by this sorting step, we can form the clusters by scanning through the sorted list once. From the clusters we obtain the discriminating regions. Subsequently, we sort the clusters based on their lengths. Finally we form the model sequence for *g*_*i*_. Across all the genomes, this can be done in one pass through the data (assuming that there is space in core memory for each genome alone).

##### Taxonomic ranks computation

The database on the model sequences can be constructed in memory. We can also compute the ranks in memory. We can align the reads in one pass through the reads.

Putting together, the algorithm takes 4 passes through the genome dataset and one pass through the read dataset. Clearly, the algorithm is asymptotically optimal in its I/O complexity.

## 3 Results

### 3.1 Datasets employed

At first we downloaded summarized information (such as, taxonomic ids, scientific names, download links, etc.) of all bacteria available in *GeneBank* from ftp.ncbi.nlm.nih.gov/genomes/genbank/bacteria/assembly_summary.txt. We then parsed the downloaded file to select a set of genomes having complete genome sequences deposited in *GeneBank*. Please note that we identify a single genome for each unique *species_taxid* to download an identical copy of the genome sequence. Where possible the *representative* genome was selected for each unique taxonomic id; however, if no *representative* genome was available, a single genome was randomly selected from all the accessions annotated as *Complete Genome*.

### 3.2 *Mock* Microbial Metagenomes

A mock community of 36 bacterial species prevalent in the human microbiome [21] was created as described by [6]. Sequencing libraries were created using the *Illumina TruSeq Nano DNA HT kit* and sequenced on the *Illumina HiSeq 2000* platform to generate 2 × 150 bp paired-end reads. Prior to analysis reads were processed with Trimmomatic to remove sequencing adapters. Following the same procedure we also generated another mock community having 11 bacterial species.

### 3.3 *In Silico* Microbial Metagenomes

We simulated 6 metagenomic paired-end *in silico* datasets by employing various Illumina platforms. Reads are produced by engaging a shotgun sequence simulator named InSilicoSeq [8]. At first we randomly selected 200 genomes from around 6k “complete” genomes from *GeneBank*. Please, note that all the randomly selected genomes also appeared in the databases of CLARK and Kraken. We generated simulated paired-end reads D1-D6 containing randomly chosen 50 and 100 bacterial species from the set of 200 genomes as stated above by employing 3 popular Illumina error models (e.g., HiSeq, MiSeq, and NovaSeq) with realistic abundance distribution. Figure III demonstrates the abundance distribution of bacterial species in each type of Illumina platforms. An example of the script for read generation is as follows: `iss generate --genomes random50.fasta --n_reads 3000000 --model hiseq --output hiseq50.fastq`. Here “random50.fasta” contains the randomly chosen 50 bacterial genomes “random50.fasta” and “hiseq50.fastq” contains the 3M simulated HiSeq reads based on the randomly chosen genomes with built-in realistic abundance distribution. The details of the datasets can be found in Table I.

**Table I.** Dataset information.

| Dataset | Model | Genomes | Paired-end reads | Read length |
| --- | --- | --- | --- | --- |
| D1 | HiSeq | 50 | 3M | 125 |
| D2 | HiSeq | 100 | 5M | 125 |
| D3 | MiSeq | 50 | 3M | 300 |
| D4 | MiSeq | 100 | 5M | 300 |
| D5 | NovaSeq | 50 | 3M | 150 |
| D6 | NovaSeq | 100 | 5M | 150 |
| D7 | HiSeq | 11 | 4.4M | 150 |
| D8 | HiSeq | 36 | 6.1M | 150 |

**Figure I.**
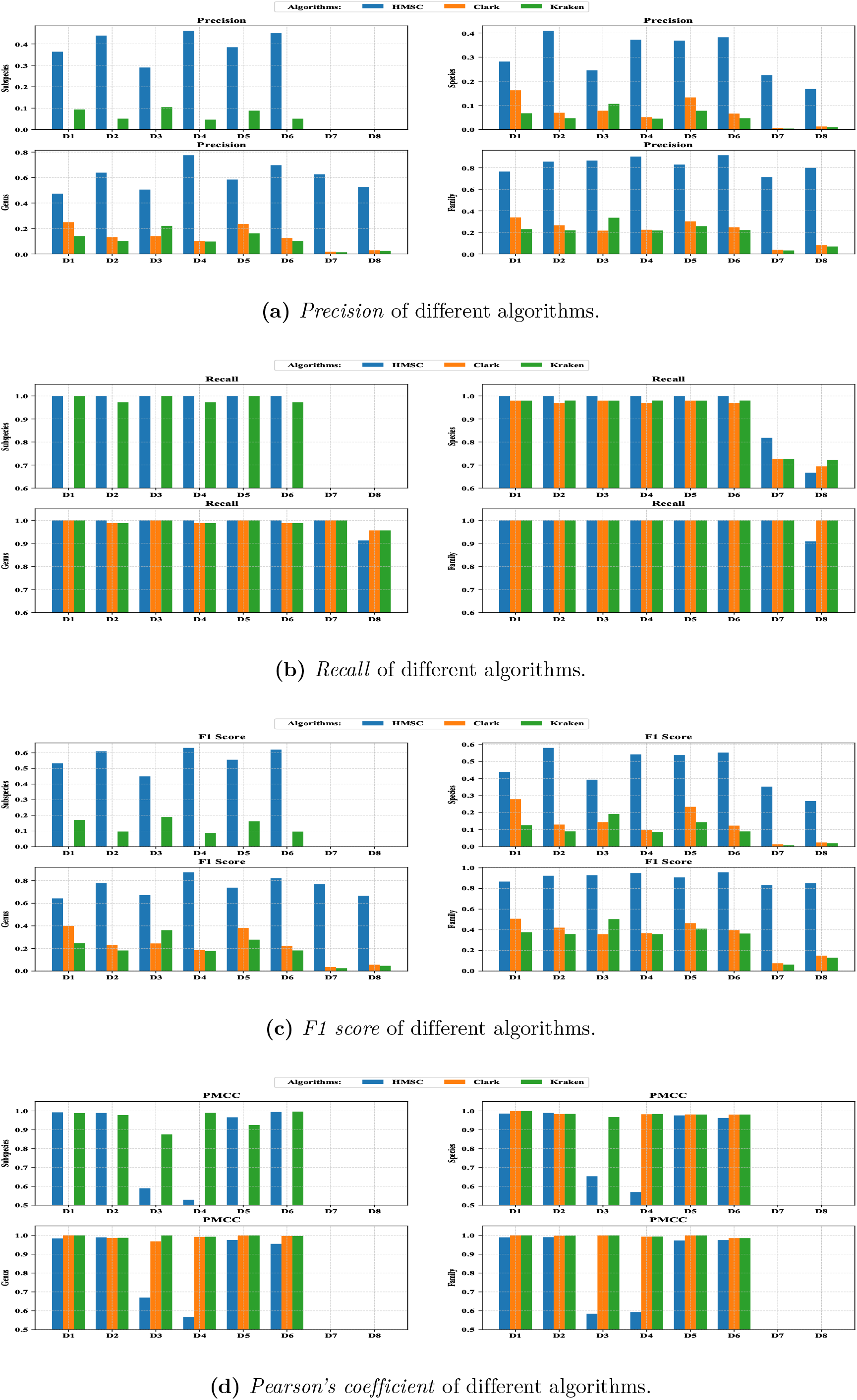
Performance evaluations.

**Figure II.**
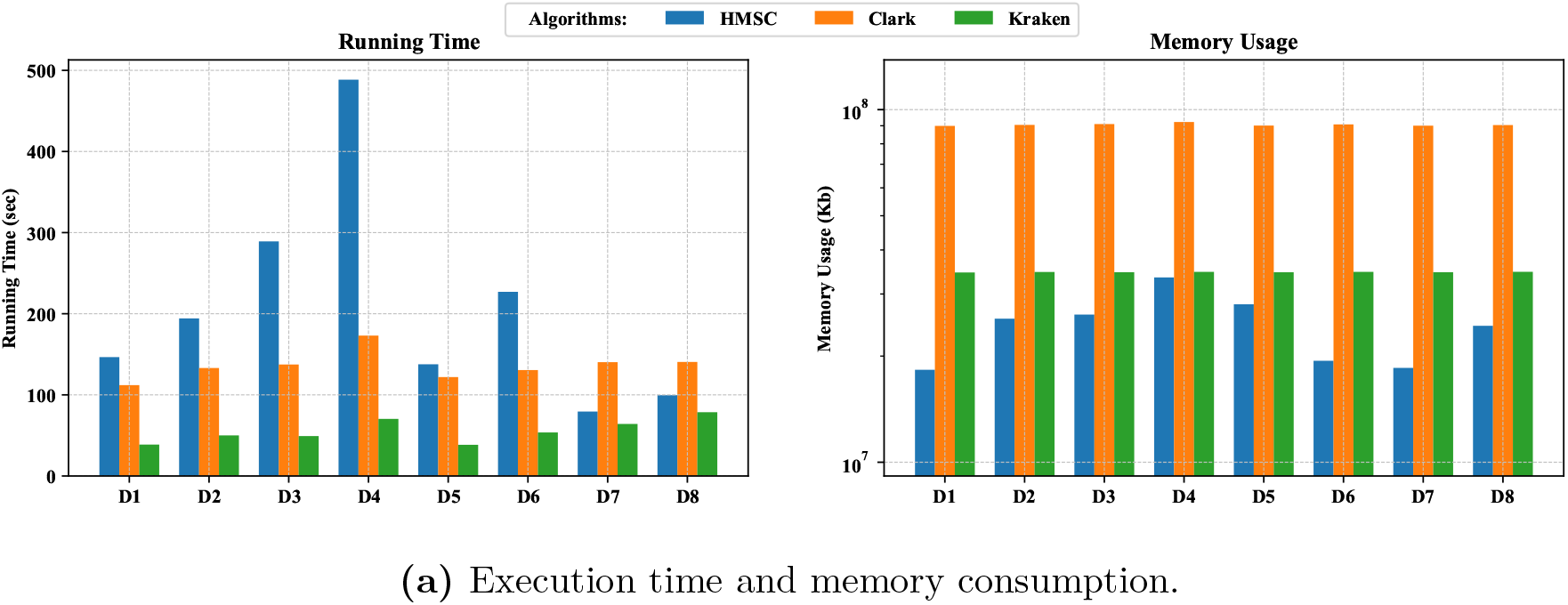
Performance evaluations.

**Figure III.**
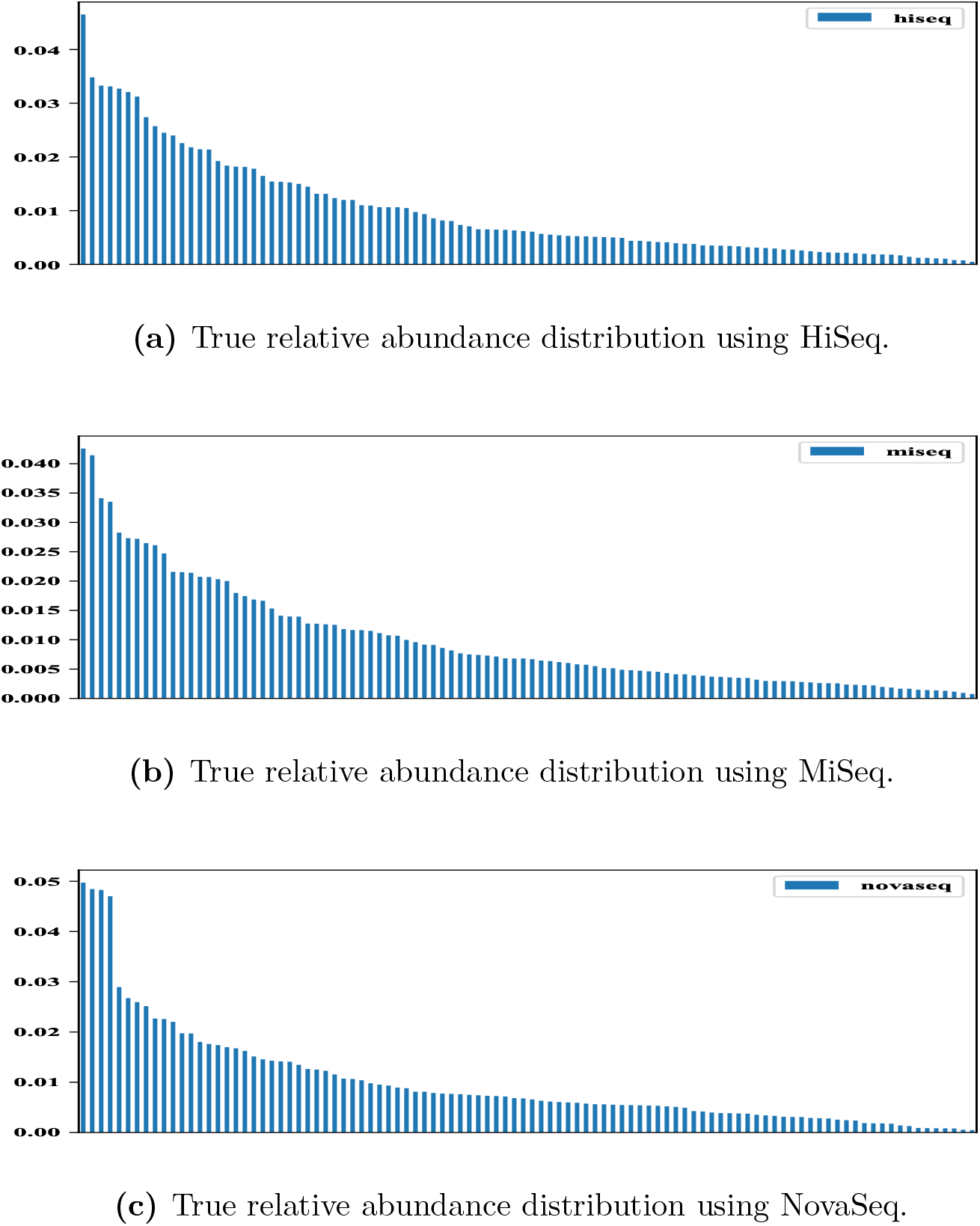
True relative abundances of 100 species.

### 3.4 Experimental setup

All the algorithms are evaluated on a Dell Precision Workstation T7910 running RHEL 7.0 on two sockets each containing 8 Dual Intel Xeon Processors E5-2667 (8C 16HT, 20MB Cache, 3.2GHz) and 256GB RAM. HMSC is written in standard Java programming language. Java source code is compiled and run by Java Virtual Machine (JVM) 1.8.0. Elapsed time was measured by taking the wall clock time.

### 3.5 Taxonomic Profiling of Microbial Metagenomes

To provide a benchmark of taxonomic abundance estimation in *in silico* (D1-D6) and *mock* (D7-D8) bacterial communities, CLARK V1.2.5.1 was downloaded and installed following online instructions (http://clark.cs.ucr.edu) and a discriminatory *k*-mer database for bacteria/archaea was created on 02/11/2018 using the default script `set_targets.sh` with the database option `bacteria`. Kraken2 was downloaded from https://ccb.jhu.edu/software/kraken2/ and a *k*-mer database for bacteria built on 02/11/2018 using the command `kraken2-build –standard –db` followed by `kraken2-build –standard –threads 10 –db`.

Taxonomic profiling with Kraken2 was performed with the command-line options `kraken2 –db <database> –threads 10 <metagenomic_file>.fastq`. Taxonomic profiling with CLARK was run using the script `classify_metagenome.sh` with the options `-O <metagenomic_file>.fastq -R <clarkoutput> -n 10`, followed by `estimate_abundance.sh` with options `-F <clarkoutput> -D <database>`.

### 3.6 Performance metrics

To demonstrate the utility of unique *k*-mer based model sequences for taxonomic profiling we compared the performance of HMSC with two widely used, *k*-mer-based tools, CLARK and Kraken, selected for their high accuracy and low execution time. We computed the following four performance metrics to demonstrate the efficacy of our algorithm HMSC:

#### Recall

In information retrieval, *recall* is the fraction of the taxa level (e.g., genus, species, etc.) in the metagenomic sample that are successfully detected. It is defined as: 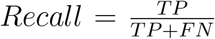. Here TP stands for the number of true positives, i.e., the number of taxa present in the sample and correctly identified by an algorithm. FN stands for the number of false negatives, i.e., the number of taxa present in the sample and not identified by an algorithm.

#### Precision

It is the fraction of retrieved taxa levels that are residing in the sample. The definition is: 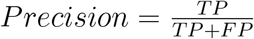. Here FP is the number of false positives, i.e., the number of taxa identified by an algorithm that are not in the sample.

#### F measure

It is the harmonic mean of *precision* and *recall*, the traditional F-measure or balanced F-score: 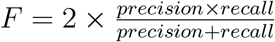. This measure is approximately the average of the two when they are close, and is more generally the harmonic mean, which, for the case of two numbers, coincides with the square of the geometric mean divided by the arithmetic mean.

#### Gain

Gain (or, improvement) measures the percentage improvement in *F measure* achieved by HMSC when compared with other algorithms. Let the *F measures* of HMSC and other algorithm of interest be *f ′* and *f ″*, respectively. The gain is measured using this formulation: 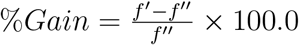.

#### Pearson’s Moment Correlation Coefficient (PMCC)

We use PMCC to measure how much estimated relative abundances of taxonomic ranks are correlated with respect to ground truths. In statistics, the Pearson moment correlation coefficient (PCC), also referred to as Pearson’s *r*, is a measure of the linear correlation between two variables *X* and *Y*. We can think of *X* and *Y* as 2 vectors each having *N* entries. According to the Cauchy–Schwarz inequality it has a value between +1 and −1, where 1 is total positive linear correlation, 0 is no linear correlation, and −1 is total negative linear correlation. The mathematics formulation is: 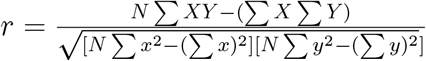.

We know the unique taxonomic id of each microbe residing in the simulated datasets *a priori*. From a taxonomic id we can retrieve all the taxonomic ranks of a microbe by traversing the taxonomy tree of life. From these ground truths (i.e, taxonomic ids) of all the datasets we identify each taxonomic rank *t* of all the microbes. Suppose *A* that belongs to a specific taxonomic rank *t* (such as, genus). For each algorithm we also identify the same taxonomic rank *t* predicted, *B*. We compute *recall* and *precision* as |*A* ∩ *B*|*/*|*A*| and |*A* ∩ *B*|*/*|*B*|, respectively. For each algorithm we compute the performance metrics for running each taxonomic profiler on the *mock* and *in silico* communities.

### 3.7 Precision, recall, and F measure

The performance of a classification algorithm depends on both *precision* and *recall*. An algorithm can have a high *recall* but a small *precision* due to the fact that the algorithm with a low *precision* suffers from high false positives. To logically fix the issue the classification performance of an algorithm is measured by taking the harmonic mean of *recall* and *precision* (known as *F measure*). It is observed that the existing algorithms for classifying metagenomic sequences suffer from very high false positives i.e., they inaccurately identifies a large number of microbes that does not belong to the metagenomic sample.

At first, consider the *in silico* datasets. As noted earlier our *in silico* datasets consist of 6 metagenomic samples (please, see D1-D6 in Table I). HMSC possesses perfect *recall* of 1.0 for every *in silico* datasets i.e., it was able to identify all the microbes (and their associated taxonomic ranks) prevalent in the samples. Although HMSC detects microbes that are not in the samples (i.e., false positives), the numbers are far smaller than CLARK or Kraken. It is evident from *precision* and *F measure* - HMSC’s *precisions* and *F measures* are higher than CLARK and Kraken for all taxonomic ranks in every datasets. Please, note that we only show 6 taxonomic ranks (e.g., subspecies, species, genus, family, order, and class) in Table II and 4 taxonomic ranks (e.g., subspecies, species, genus, and family) in Figure I because of space constraints. To estimate HMSC’s improvement over other algorithms with respect to classification accuracy (i.e., *F measure*), we define a performance metric *Gain* as stated earlier. Table IV shows the % improvement of HMSC’s *F measure* over CLARK and Kraken. Please, note that CLARK can not predict the lowest taxonomic rank (i.e., subspecies) and hence we omitted subspecies level *Gain* computation.

**Table II.** Performance evaluations on *in silico* datasets

|  |  | HMSC |  |  |  | CLARK |  |  |  | Kraken |  |  |  |
| --- | --- | --- | --- | --- | --- | --- | --- | --- | --- | --- | --- | --- | --- |
|  |  | Recall | Precision | F-score | PMCC | Recall | Precision | F-score | PMCC | Recall | Precision | F-score | PMCC |
| D1 | Subspecies | 1.0000 | 0.3636 | 0.5333 | 0.9924 | NA | NA | NA | NA | 1.0000 | 0.0935 | 0.1709 | 0.9884 |
|  | Species | 1.0000 | 0.2816 | 0.4395 | 0.9860 | 0.9796 | 0.1627 | 0.2791 | 0.9998 | 0.9796 | 0.0673 | 0.1260 | 0.9998 |
|  | Genus | 1.0000 | 0.4737 | 0.6429 | 0.9839 | 1.0000 | 0.2500 | 0.4000 | 1.0000 | 1.0000 | 0.1402 | 0.2459 | 0.9999 |
|  | Family | 1.0000 | 0.7647 | 0.8667 | 0.9896 | 1.0000 | 0.3391 | 0.5065 | 1.0000 | 1.0000 | 0.2308 | 0.3750 | 1.0000 |
|  | Order | 1.0000 | 0.7500 | 0.8571 | 0.9949 | 1.0000 | 0.4167 | 0.5882 | 1.0000 | 1.0000 | 0.3333 | 0.5000 | 1.0000 |
|  | Class | 1.0000 | 0.7200 | 0.8372 | 0.9996 | 1.0000 | 0.5625 | 0.7200 | 1.0000 | 1.0000 | 0.4286 | 0.6000 | 1.0000 |
| D2 | Subspecies | 1.0000 | 0.4390 | 0.6102 | 0.9892 | NA | NA | NA | NA | 0.9722 | 0.0508 | 0.0966 | 0.9776 |
|  | Species | 1.0000 | 0.4091 | 0.5806 | 0.9897 | 0.9697 | 0.0695 | 0.1297 | 0.9835 | 0.9798 | 0.0470 | 0.0897 | 0.9846 |
|  | Genus | 1.0000 | 0.6385 | 0.7793 | 0.9895 | 0.9880 | 0.1312 | 0.2316 | 0.9861 | 0.9880 | 0.1004 | 0.1822 | 0.9870 |
|  | Family | 1.0000 | 0.8571 | 0.9231 | 0.9903 | 1.0000 | 0.2661 | 0.4204 | 0.9979 | 1.0000 | 0.2185 | 0.3587 | 0.9985 |
|  | Order | 1.0000 | 0.8367 | 0.9111 | 0.9950 | 1.0000 | 0.3361 | 0.5031 | 0.9987 | 1.0000 | 0.2971 | 0.4581 | 0.9989 |
|  | Class | 1.0000 | 0.8261 | 0.9048 | 0.9990 | 1.0000 | 0.3276 | 0.4935 | 0.9998 | 1.0000 | 0.3016 | 0.4634 | 0.9997 |
| D3 | Subspecies | 1.0000 | 0.2899 | 0.4494 | 0.5898 | NA | NA | NA | NA | 1.0000 | 0.1047 | 0.1896 | 0.8758 |
|  | Species | 1.0000 | 0.2450 | 0.3936 | 0.6536 | 0.9796 | 0.0779 | 0.1444 | 96.11 | 0.9796 | 0.1064 | 0.1920 | 0.9670 |
|  | Genus | 1.0000 | 0.5056 | 0.6716 | 0.6697 | 1.0000 | 0.1393 | 0.2446 | 0.9682 | 1.0000 | 0.2206 | 0.3614 | 0.9995 |
|  | Family | 1.0000 | 0.8667 | 0.9286 | 0.5844 | 1.0000 | 0.2167 | 0.3562 | 0.9995 | 1.0000 | 0.3362 | 0.5032 | 0.9996 |
|  | Order | 1.0000 | 0.9091 | 0.9524 | 0.7992 | 1.0000 | 0.3061 | 0.4687 | 0.9995 | 1.0000 | 0.4688 | 0.6383 | 0.9997 |
|  | Class | 1.0000 | 0.9000 | 0.9474 | 0.8471 | 1.0000 | 0.3913 | 0.5625 | 0.9997 | 1.0000 | 0.6667 | 0.8000 | 0.9998 |
| D4 | Subspecies | 1.0000 | 0.4615 | 0.6316 | 0.5287 | NA | NA | NA | NA | 0.9722 | 0.0461 | 0.0879 | 0.9902 |
|  | Species | 1.0000 | 0.3722 | 0.5425 | 0.5698 | 0.9697 | 0.0515 | 0.0979 | 0.9823 | 0.9798 | 0.0450 | 0.0861 | 0.9835 |
|  | Genus | 1.0000 | 0.7757 | 0.8737 | 0.5672 | 0.9880 | 0.1026 | 0.1859 | 0.9918 | 0.9880 | 0.0976 | 0.1777 | 0.9926 |
|  | Family | 1.0000 | 0.9041 | 0.9496 | 0.5936 | 1.0000 | 0.2245 | 0.3667 | 0.9932 | 1.0000 | 0.2178 | 0.3577 | 0.9938 |
|  | Order | 1.0000 | 0.9535 | 0.9762 | 0.6932 | 1.0000 | 0.3015 | 0.4633 | 0.9956 | 1.0000 | 0.2847 | 0.4432 | 0.9958 |
|  | Class | 1.0000 | 0.9500 | 0.9744 | 0.8718 | 1.0000 | 0.2923 | 0.4524 | 0.9986 | 1.0000 | 0.2836 | 0.4419 | 0.9983 |
| D5 | Subspecies | 1.0000 | 0.3846 | 0.5556 | 0.9661 | NA | NA | NA | NA | 1.0000 | 0.0881 | 0.1619 | 0.9251 |
|  | Species | 1.0000 | 0.3684 | 0.5385 | 0.9761 | 0.9796 | 0.1330 | 0.2341 | 0.9813 | 0.9796 | 0.0777 | 0.1439 | 0.9810 |
|  | Genus | 1.0000 | 0.5844 | 0.7377 | 0.9754 | 1.0000 | 0.2356 | 0.3814 | 0.9993 | 1.0000 | 0.1613 | 0.2778 | 0.9995 |
|  | Family | 1.0000 | 0.8298 | 0.9070 | 0.9728 | 1.0000 | 0.3023 | 0.4643 | 0.9994 | 1.0000 | 0.2583 | 0.4105 | 0.9996 |
|  | Order | 1.0000 | 0.8571 | 0.9231 | 0.9745 | 1.0000 | 0.4167 | 0.5882 | 0.9995 | 1.0000 | 0.3371 | 0.5042 | 0.9996 |
|  | Class | 1.0000 | 0.8571 | 0.9231 | 0.9799 | 1.0000 | 0.6000 | 0.7500 | 0.9997 | 1.0000 | 0.4615 | 0.6316 | 0.9998 |
| D6 | Subspecies | 1.0000 | 0.4500 | 0.6207 | 0.9948 | NA | NA | NA | NA | 0.9722 | 0.0507 | 0.0963 | 0.9966 |
|  | Species | 1.0000 | 0.3822 | 0.5531 | 0.9624 | 0.9697 | 0.0659 | 0.1235 | 0.9807 | 0.9798 | 0.0469 | 0.0896 | 0.9809 |
|  | Genus | 1.0000 | 0.6975 | 0.8218 | 0.9551 | 0.9880 | 0.1252 | 0.2222 | 0.9967 | 0.9880 | 0.1006 | 0.1826 | 0.9968 |
|  | Family | 1.0000 | 0.9167 | 0.9565 | 0.9751 | 1.0000 | 0.2472 | 0.3964 | 0.9854 | 1.0000 | 0.2222 | 0.3636 | 0.9855 |
|  | Order | 1.0000 | 0.9318 | 0.9647 | 0.9890 | 1.0000 | 0.3083 | 0.4713 | 0.9941 | 1.0000 | 0.2971 | 0.4581 | 0.9941 |
|  | Class | 1.0000 | 0.9048 | 0.9500 | 0.9935 | 1.0000 | 0.3065 | 0.4691 | 0.9967 | 1.0000 | 0.2923 | 0.4524 | 0.9966 |

**Table III.** Performance evaluations on *mock* datasets.

|  |  | HMSC |  |  | CLARK |  |  | Kraken |  |  |
| --- | --- | --- | --- | --- | --- | --- | --- | --- | --- | --- |
|  |  | Recall | Precision | F1 score | Recall | Precision | F1 score | Recall | Precision | F1 score |
| **D7** | **Species** | 0.8182 | 0.2250 | 0.3529 | 0.7273 | 0.0069 | 0.0138 | 0.7273 | 0.0040 | 0.0080 |
|  | **Genus** | 1.0000 | 0.6250 | 0.7692 | 1.0000 | 0.0181 | 0.0357 | 1.0000 | 0.0129 | 0.0254 |
|  | **Family** | 1.0000 | 0.7143 | 0.8333 | 1.0000 | 0.0394 | 0.0758 | 1.0000 | 0.0322 | 0.0623 |
|  | **Order** | 1.0000 | 0.7273 | 0.8421 | 1.0000 | 0.0625 | 0.1176 | 1.0000 | 0.0544 | 0.1032 |
|  | **Class** | 1.0000 | 0.6000 | 0.7500 | 1.0000 | 0.0984 | 0.1791 | 1.0000 | 0.0870 | 0.1600 |
| **D8** | **Species** | 0.6667 | 0.1678 | 0.2682 | 0.6944 | 0.0126 | 0.0248 | 0.7222 | 0.0100 | 0.0198 |
|  | **Genus** | 0.9130 | 0.5250 | 0.6667 | 0.9565 | 0.0288 | 0.0558 | 0.9565 | 0.0238 | 0.0464 |
|  | **Family** | 0.9091 | 0.8000 | 0.8511 | 1.0000 | 0.0806 | 0.1492 | 1.0000 | 0.0690 | 0.1290 |
|  | **Order** | 0.9375 | 0.8824 | 0.9091 | 1.0000 | 0.1290 | 0.2286 | 1.0000 | 0.1067 | 0.1928 |
|  | **Class** | 0.9000 | 0.9000 | 0.9000 | 1.0000 | 0.1724 | 0.2941 | 1.0000 | 0.1408 | 0.2469 |

**Table IV.** % Improvement over CLARK and Kraken.

|  |  | CLARK |  |  |  | Kraken |  |  |  |
| --- | --- | --- | --- | --- | --- | --- | --- | --- | --- |
|  |  | Species | Genus | Family | Order | Species | Genus | Family | Order |
| D1 |  | 57.47 | 60.73 | 71.12 | 45.72 | 248.81 | 161.45 | 131.12 | 71.42 |
| D2 |  | 347.65 | 236.49 | 119.58 | 81.10 | 547.27 | 327.72 | 157.35 | 98.89 |
| D3 |  | 172.58 | 174.57 | 160.70 | 103.20 | 105.00 | 85.83 | 84.54 | 49.21 |
| D4 |  | 454.14 | 369.98 | 158.96 | 110.71 | 530.08 | 391.67 | 165.47 | 120.26 |
| D5 |  | 130.03 | 93.42 | 95.35 | 56.94 | 274.22 | 165.55 | 120.95 | 83.08 |
| D6 |  | 347.85 | 269.85 | 141.30 | 104.69 | 517.30 | 350.05 | 163.06 | 110.59 |
| D7 |  | 2,457.25 | 2,054.62 | 999.34 | 616.07 | 4,311.25 | 2,928.35 | 1,237.56 | 715.99 |
| D8 |  | 981.45 | 1,094.80 | 470.44 | 297.68 | 1,254.55 | 1,336.85 | 559.77 | 371.52 |

Now consider mock datasets (D7-D8). In D7 dataset HMSC’s *recall* of species-level taxonomy is better than that of CLARK and Kraken. In all other cases *recall* measures are identical for all 3 algorithms. On the contrary *precision* and *F* measures are higher than that of CLARK and Kraken in both of D7 and D8 datasets. To get an estimate of % improvement of HMSC’s *F measures* over CLARK and Kraken please, see Table IV. It is evident from the table that both of the algorithms erroneously identify a lot of microbes that are not residing in the sample.

### 3.8 Relative abundances

Our algorithm HMSC deals with the model sequences of the reference genomes. Each model sequence comprises a set of discriminating regions of a genome. Therefore, all the reads coming from a specific genome will not be aligned onto the model sequence designated for that genome. Only the discriminating reads will be aligned onto a specific model sequence. As model sequences contains discriminating stretches of genome sequences, it estimates approximate relative abundances instead of true relative abundances. Although in theory the approximation may be over-represented or under-represented, HMSC mimics, in practice, the true relative abundances in most of cases. Since we do not know the true relative abundances of the 2 mock communities (e.g., D7-D8), we could not be able to compute Pearson’s correlations. Please, see Figure I[d] for visual comparisons of different methods employed including HMSC. It is to be noted that the abundance estimations of HMSC in MiSeq datasets are poor with respect to CLARK and Kraken. It might be due to the fact that we are aligning reads onto model sequences without any mismatches for every datasets to preserve uniformity. Since the length of MiSeq reads is 300bp long, many of them might not be aligned onto the model sequences within the mismatch threshold we used (i.e., *d* = 0). Currently, we are investigating this issue.

### 3.9 Execution time and memory consumption

HMSC has a very low memory footprint. On average, HMSC uses 3.70× and 1.43× less memory than that of CLARK and Kraken, respectively. The execution times of HMSC are also comparable with the state-of-the-art algorithms. In general, HMSC is faster than CLARK on real datasets D7 and D8. Comparing to Kraken, HMSC’s running time is in the same order of magnitude. Please, see Figure II for visual comparisons.

## 4 Discussion

The algorithmic approach followed by HMSC is fundamentally different from the other techniques in the domain of metagenomic sequence classification. It is designed to answer the most important question with a high level of confidence: What are those microbioms interacting in a complex and diversified metagenomic sample? To answer this question accurately we search through the reference genomes, extract unique *k*-mers, find discriminating regions, and build model sequences for each genome. We call our algorithm “hybrid” as it exploits both alignment-free and alignment-based techniques. Please, see Methods section for more details about the approaches stated above. At the first stage HMSC explores entire space of reference genomes to find unique *k*-mers that is essentially an alignment-free approach. Next unique *k*-mers are clustered to extract discriminating regions. By employing those regions HMSC then builds a model sequence for each reference genome. Metagenomic reads are then aligned onto those model sequences which is actually an alignment-based approach. This is why we call our algorithm “hybrid.”

As discussed in Methods section each model sequence of a particular reference genome contains a subset of consecutive unique *k*-mer that does not belong to any other reference genomes. If a metagenomic read is aligned onto a model sequence, it must contain at least one of the unique *k*-mers depending on the read length. For an example let the distance between 2 consecutive *k*-mers be at most *l′* and the read length be *l″*. If *l″* ≥ 2 × *k* + *l′*, it is guaranteed that the read must contains at least one unique *k*-mer. Since in our experiment we set *k* = 31 and *l′* = 40, a read with length at least 102 (a reasonable assumption) must entirely contains at least one of the unique *k*-mers given that the read is aligned onto the model sequence. If *l″ <* 2 × *k* + *l′*, it may or may not totally contain a unique *k*-mer. However the read must partially contains one or more unique *k*-mers given that the precondition *l″* ≥ *l′* satisfies. All the above assumptions we have made are based on worst case scenario. In practice distance between consecutive pairs of unique *k*-mers may be well below the threshold *l′* and so, an aligned read can contain multiple unique *k*-mers. Moreover the longer the read the higher the number of unique *k*-mers it will contain given that the read is aligned onto a model sequence. So, if a read is aligned onto a model sequence, it will be highly discriminating in classifying a microbe present in the sample.

Note that in alignment-based approach a metagenomic sequence read can be aligned onto multiple reference genomes within certain mismatches (e.g. insertions, deletions, and/or substitutions). Mismatches can be occurred because of purely biological events (such as, mutations, recombinations, etc.) or limitations of the sequencing techniques used. In this case we don’t have much choices but to classify that reads for all the genomes it aligned onto. It also applies to alignment-free approach where a unique *k*-mer can be found into multiple reads within certain mismatches due to the identical facts stated above. In our algorithm a model sequence contains a set of highly discriminating regions comprising a set of unique *k*-mers. If a read aligned onto a model sequence, it not only contains unique *k*-mer(s), but also surrounding nucleotides of a particular genome. As a result the metagenomic reads aligned onto the model sequences are also highly discriminating in nature. Based on these reads we classify the taxonomic ranks of each microbe present in a metagenomic sample with a high level of confidence. This is why our experimental evaluations show that HMSC outputs very few false positives compared to the other state-of-the-art algorithms. Please, note that unlike CLARK, HMSC can detect the lowest taxonomic ranks (i.e., subspecies).

It is also noteworthy that both Kraken and CLARK dramatically over-predicted the number of taxonomic ranks present in each community. While false positive taxa were generally detected at low relative abundance, their presence may still affect biological interpretation of taxonomic profiling results. In addition to the biological relevance of low abundance taxa [10], their prevalence can influence commonly used *alpha* diversity measures such as *richness* and *chao1* [4]. They will also have a bearing on statistical approaches that attempt to accommodate for zero-inflation in microbiome datasets [27]. Compared to the other tools HMSC had far fewer incorrect predictions in all the taxonomic levels in average, suggesting that appropriate selection of the discriminatory regions has the potential to address the problem of false positive taxonomic ranks in metagenomic sequencing datasets.

In addition, another question has to be answered: What are the relative abundances among the taxonomic ranks? The total size of all the reference genomes is 21.5GB where the model sequences consume only 4.5GB disk space - approximately 5× reduction. We align metagenomic sequencing reads onto the model sequences and identify all taxonomic ranks. As HMSC deals with only a small part of the reference genomic sequences, it estimates relative abundances approximately. In theory it may not reflect the true abundances prevalent in the metagenomic sample. But in practice it is visible from the experimental evaluations that HMSC indeed excels in estimating the relative abundances of microbes present in the sample.

## 5 Conclusion

In this article we propose HMSC that can accurately detect microbes and their relative abundances in a metagenomic sample. The algorithm judiciously exploits both alignment-free and alignment-based approaches and our rigorous experimental evaluations show that it is indeed an effective, scalable, and efficient algorithm compared to the other state-of-the-art methods in terms of accuracy, memory, and runtime.

## Algorithm 1 Mining Unique K-mers (MUK)

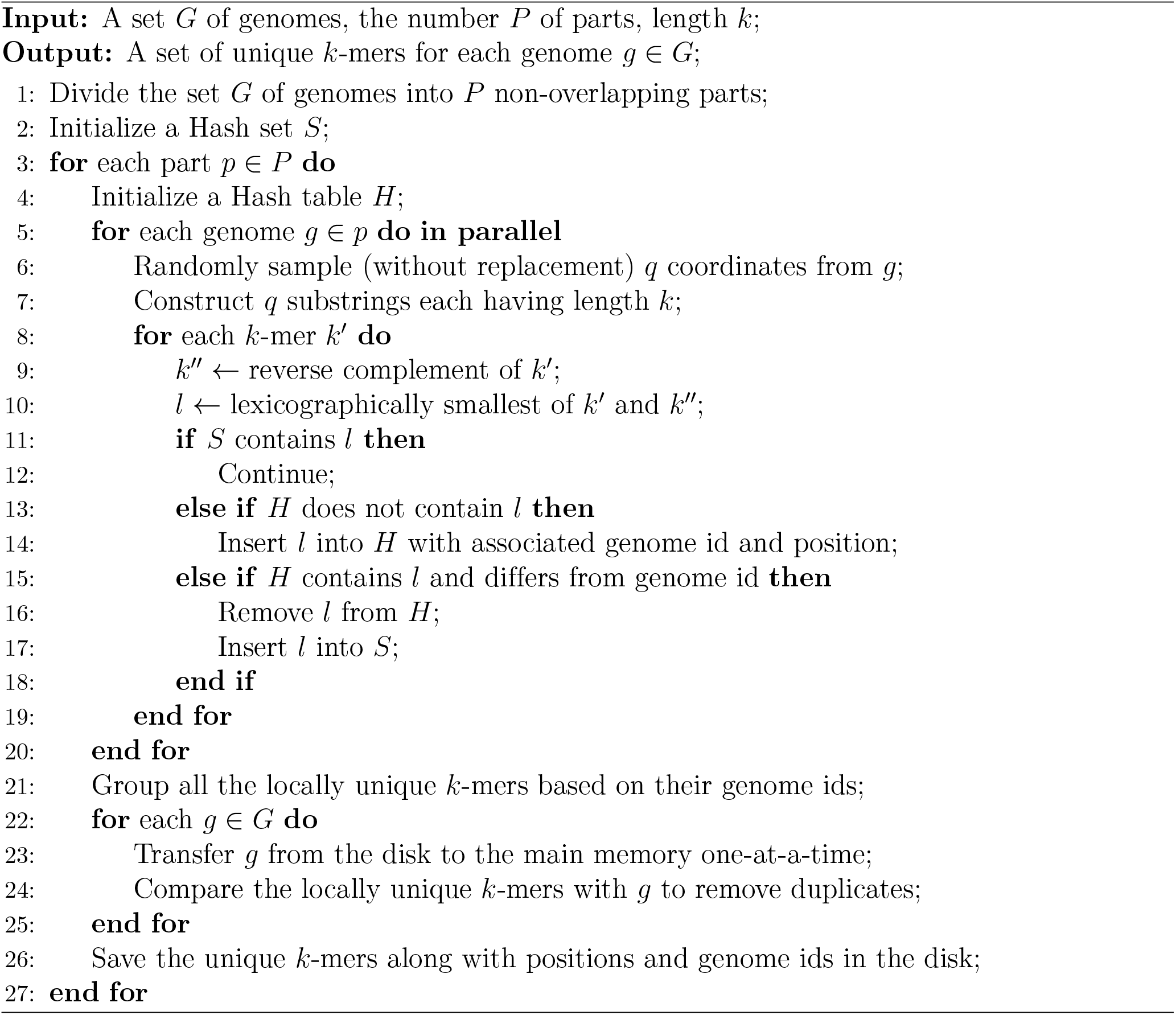

## Algorithm 2 Building Model Sequences (BMS)

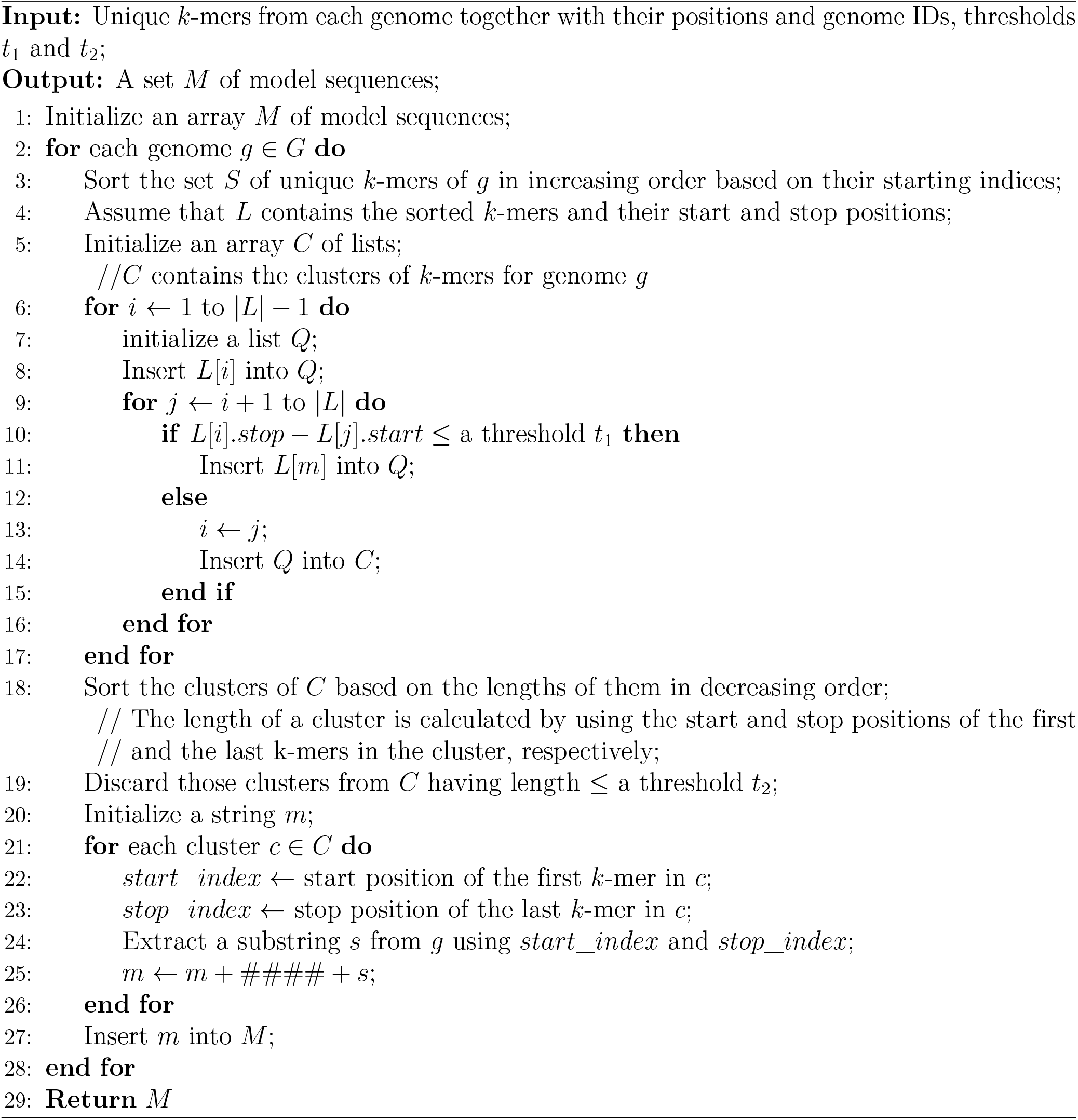

